# The role of the cerebellum in learning to predict reward: evidence from cerebellar ataxia

**DOI:** 10.1101/2022.11.04.515251

**Authors:** Jonathan Nicholas, Christian Amlang, Chi-Ying R. Lin, Leila Montaser-Kouhsari, Natasha Desai, Ming-Kai Pan, Sheng-Han Kuo, Daphna Shohamy

## Abstract

Recent findings in animals have challenged the traditional view of the cerebellum solely as the site of motor control, suggesting that the cerebellum may also be important for learning to predict reward from trial-and-error feedback. Yet, evidence for the role of the cerebellum in reward learning in humans is lacking. Moreover, open questions remain about which specific aspects of reward learning the cerebellum may contribute to. Here we address this gap through an investigation of multiple forms of reward learning in individuals with cerebellum dysfunction, represented by cerebellar ataxia cases. Nineteen participants with cerebellar ataxia and 57 age- and sex-matched healthy controls completed two separate tasks that required learning about reward contingencies from trial-and-error. To probe the selectivity of reward learning processes, the tasks differed in their underlying structure: while one task measured incremental reward learning ability alone, the other allowed participants to use an alternative learning strategy based on episodic memory alongside incremental reward learning. We found that individuals with cerebellar ataxia were profoundly impaired at reward learning from trial-and-error feedback on both tasks, but retained the ability to learn to predict reward based on episodic memory. These findings provide evidence from humans for a specific and necessary role for the cerebellum in incremental learning of reward associations based on reinforcement. More broadly, the findings suggest that alongside its role in motor learning, the cerebellum likely operates in concert with the basal ganglia to support reinforcement learning from reward.

## Introduction

It is well established that the cerebellum is required for refining movement through supervised motor learning.^1–4^ The cerebellum receives error signals from climbing fiber input which then alters Purkinje cell plasticity to adapt motor behavior in service of minimizing future error.^5–7^ However, recent findings have challenged the notion that the cerebellum is solely responsible for supervised learning of motor behavior and instead suggest that the cerebellum may also be involved in the processing of reward more generally^8–19^. In particular, climbing fiber inputs to the cerebellum encode expected reward^13, 15, 17, 19^, and cerebellar Purkinje cells have been found to report reward-based prediction errors^11, 12, 18^. These signals are essential ingredients for reinforcement learning, or learning that allows an organism to determine from trial-and-error feedback which actions should be taken in order to maximize future expected reward. The presence of reward-related processing in the cerebellum suggests that it may play a role in reinforcement learning alongside its capacity for supervised motor learning^20^. This proposal challenges not only our current understanding of cerebellar function, but also our understanding of how the brain learns from reward more broadly^5, 21^.

Although research on the cerebellum’s function in reward learning is growing, the vast majority of work has been done in animal models^10–19^, and evidence in humans remains limited. Human neuroimaging studies have revealed correlational evidence that the cerebellum is involved in tasks unrelated to movement^22^, however, despite some reports of BOLD activity in the cerebellum in response to reward across several early imaging studies^23–25^, more direct investigations of the role of cerebellum in reward-related behaviors in humans are lacking. The aim of the present study was to fill this gap by testing whether individuals with damage to the cerebellum, as occurs in cerebellar ataxia (CA), are impaired in their ability to acquire stimulus-reward associations.

Our study builds upon a rich literature focused on learning about reward from trial-and-error feedback. This process has been studied extensively using models of incremental learning, which rely on error-driven rules that summarize experiences with a running average^26–28^. During reward learning of this type, an agent uses the outcome of a recent decision to associate some stimulus with an action. Following successful learning, actions that are more likely to be rewarded are more likely to be repeated. This simple mechanism has been evoked to explain conditioning behavior and is well-captured by reward prediction error signals in midbrain dopamine neurons that project to the striatum^27, 29^. This error signal is also precisely what has been implicated in recent animal models of cerebellar contributions to reward learning^9^, suggesting an additional, albeit unclear, role for the cerebellum in this process. Whether these cerebellar contributions are actually needed for successful incremental reward learning in humans is at present unknown.

To answer this question, we asked individuals with CA to complete a series of tasks that required them to learn associations between stimuli from trial-and-error feedback in order to maximize expected reward. CA is defined as a lack of coordination caused by disorders that affect cerebellar function^30^. A large variety of conditions can cause CA, ranging from immune-mediated disease to genetic and neurodegenerative disorders. Given the presence of cerebellar dysfunction in CA cases, studying individuals with CA is a common method used to investigate the necessary physiological functions of the cerebellum in humans.

Nineteen individuals with CA and 57 age- and sex-matched healthy controls (HC) completed two tasks (**Figure 1**). The first, referred to throughout as the *incremental learning* task, allowed us to measure each participants’ ability to learn about reward incrementally. This task was motivated by recent work using a similar simplified paradigm to investigate cerebellar-based incremental learning in non-human primates^10, 11^. The second task, referred to throughout as the *multiple learning strategies* task, allowed us to measure whether any impairments were specific to incremental learning alone. In the multiple learning strategies task, learning about reward can be supported by an alternative strategy based on episodic memory for trial-unique past outcomes. Healthy adults readily use of both of these strategies in this task^31, 32^. We hypothesized that cerebellar dysfunction would lead specifically to impaired incremental reward learning relative to healthy controls.

**Figure 1.**
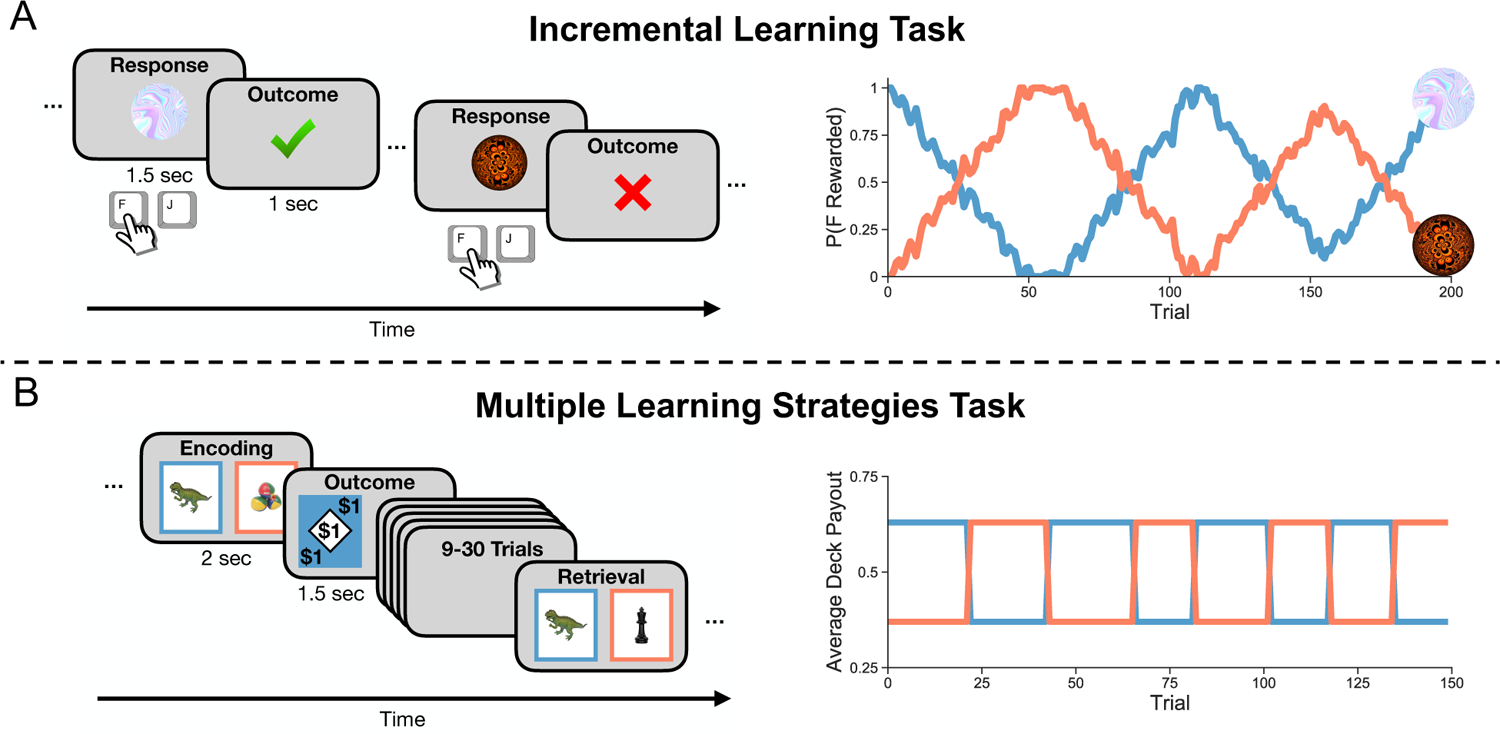
Design of the incremental learning and multiple learning strategies tasks. **(A)** Left: Trial design for the incremental learning task. Participants saw one of two fractal cues on the screen and were required to press either the F key with their left hand or the J key with their right hand. Following their choice, they received binary probabilistic feedback about whether they were correct or not. Right: Drifting cue-response-reward contingencies over the course of the incremental learning task. The probability that the F key is rewarded is shown for each cue in blue and orange. **(B)** Left: Trial design or the multiple learning strategies task. Participants chose between two decks of cards (one blue and one orange) and received an outcome between $0-$1 in intervals of 20 cents. Each card featured a trial-unique object that could repeat once every 9-30 trials. Participants were told that if they saw the same card again, it would be worth the same amount as the first time that it appeared. Right: An example of how average deck value reversed throughout the course of the multiple learning strategies task.

## Results

### Impaired reward learning in the incremental learning task

Our first goal was to assess CA participants’ baseline ability to learn incrementally from reward using the incremental learning task. On this task, CA participants made overall fewer correct choices compared to healthy controls (β*_Group_* = −0.88, 95% *CI* = [−1.55, −0.144]; **Figure 2A**). CA participants’ choices were less correct throughout the entirety of the task, even during periods of learning where action-outcome contingencies were more deterministic (e.g. close to 100%) compared to more difficult periods of learning (β*_GroupxpFReward1_^2^* = −5.49, 95% *CI* = [−7.57, −3.52]; **Figure 2B-C**. Overall, this difference in performance indicates that CA participants did not learn from reward feedback. Although CA participants responded slightly more slowly than healthy controls on this task (β*_Group_* = −115.81, 95% *CI* = [−201.26, −33.55]), we included reaction times as a covariate in the above regression analysis to ensure that differences in choice accuracy were not attributed to motor slowing in CA participants.

**Figure 2.**
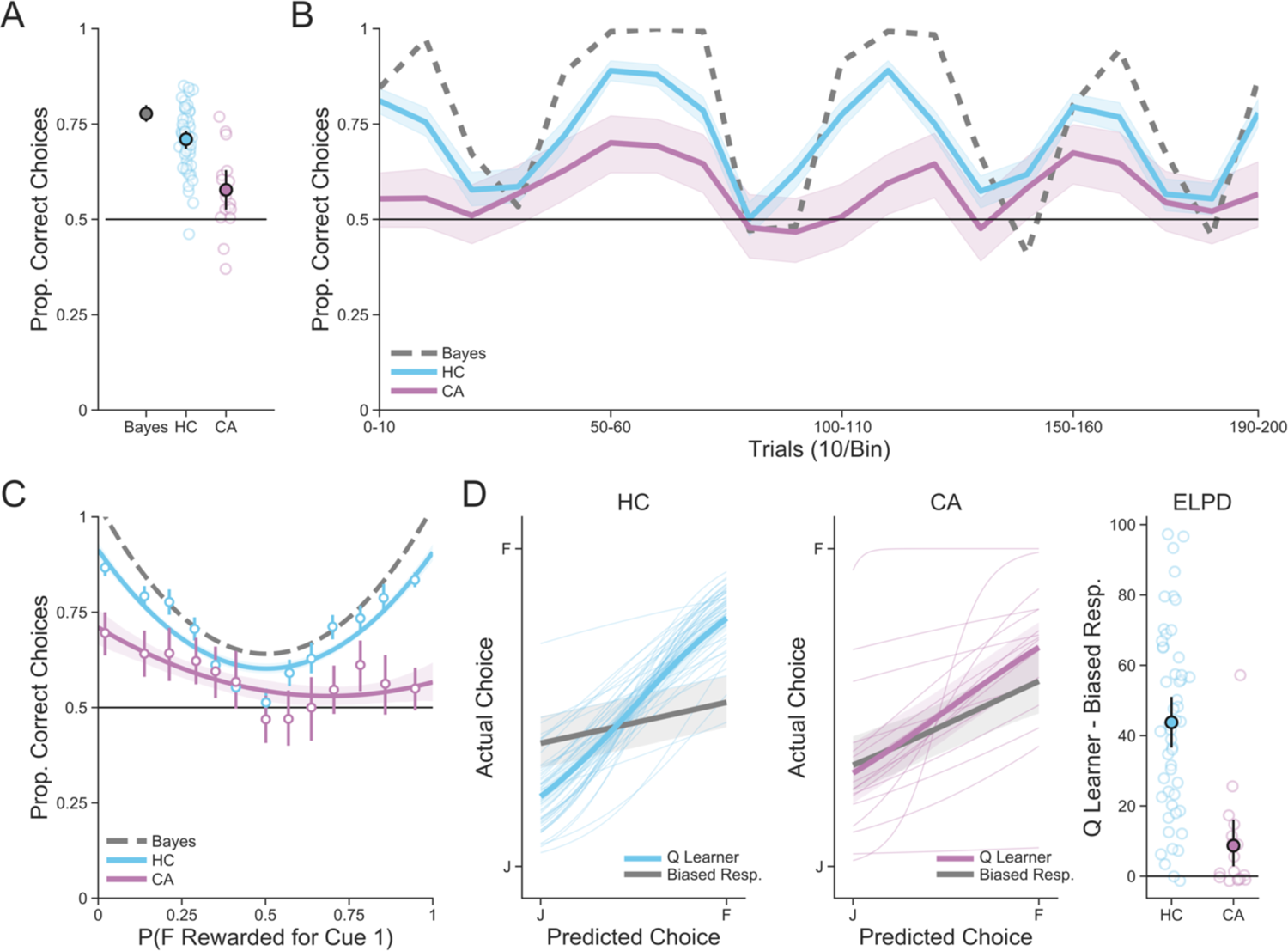
Performance on the incremental learning task. **(A)** Performance on the incremental learning task averaged across all trials for healthy controls (HC) and CA participants compared to a Bayesian observer in gray, which represents normative performance on the task. Individual points are averages for each subject and filled in points represent group-level averages. Error bars are 95% confidence intervals. **(B)** Performance on the incremental learning task over time. Each timepoint represents ten trials. Lines are group averages and bands are 95% confidence intervals. For normative comparison, the performance of the Bayesian observer is shown as a dotted gray line. **(C)** Performance on the incremental learning task as a function of task difficulty, which is indexed by the true underlying probability that pressing the F key was the correct response (>50%) on each trial. Points represent group level averages from 13 bins with an equal number of trials, lines represent the fit of a second-order linear model, and error bars and bands represent 95% confidence intervals. **(D)** Model performance of the Q Learner and baseline Biased Responder models. Left: Posterior predictive performance. Individual lines represent Q learner fits for each individual, whereas thick lines represent the group-level average fit (with the Q Learner in color and Biased Responder in gray). Bands represent 95% confidence intervals. Right: The difference in estimated out-of-sample predictive performance (as measured by expected log pointwise predictive density; ELPD) between the Q Learner and the Biased Responder model for each group. Individual points are the ELPD difference for each subject and filled in points represent group-level averages. Error bars are 95% confidence intervals.

Next, to more formally assess participants’ performance on this task, we fit a standard Q learning model to participants’ responses. This model captures the extent to which each participant incorporated trial-by-trial outcomes into running estimates of the value of pressing each button in response to each cue, as well as whether choices are based on these estimates. As a baseline, we compared the performance of this model to a biased responder that merely estimated the extent to which each participant pressed one button over the other, regardless of outcome, in response to each cue. While healthy controls’ responses were well described by the Q learning model, this model did no better than the biased responder at predicting CA participants’ decisions, thus demonstrating that CA participants engaged in little-to-no incremental learning (**Figure 2D**). On a measure of estimated out-of-sample predictive performance, controls were substantially better fit by the Q learner compared to the biased responder, while this improvement in fit was largely absent for CA participants (β_*Group*_ = 30.94, 95% *CI* = [16.465, 46.0]). Thus, while healthy controls incorporated feedback into their estimates about the relationship between cue and action at each timepoint, CA participants generally did not.

Together, these results indicate that individuals with CA are impaired at reward learning from trial-and-error.

### Impaired incremental reward learning but intact episodic memory in the multiple learning strategies task

After establishing that CA participants were impaired in a task that measured solely incremental reward learning, we wanted to examine both the specificity and generalizability of this impairment by i) providing an alternative means of reward-based decision making alongside incremental learning and ii) altering the incremental learning task structure to measure responses to reversal events rather than drifting probabilities. The multiple learning strategies task was thus used to accomplish both of these goals.

Consistent with the results of the incremental learning task, CA participants in the multiple learning strategies task were less responsive to reward outcomes compared to controls (**Figure 3A**). Specifically, controls tended to choose the lucky deck more than CA participants immediately prior to a reversal (β_*Group*×./.0-_ = 0.397, 95% *CI* = [0.002, 0.807]), and this tendency was disrupted by reversals; CA participants did not show this pattern (β_*Group*×./1_ = −0.897, 95% *CI* = [−1.28, −0.535]), and remained below chance performance after a reversal occurred (β_*Group*×./.2-_ = −0.595, 95% *CI* = [−0.984, −0.21]). This indicates that CA participants were unable to learn which deck had the higher expected value at any given time throughout the task.

**Figure 3.**
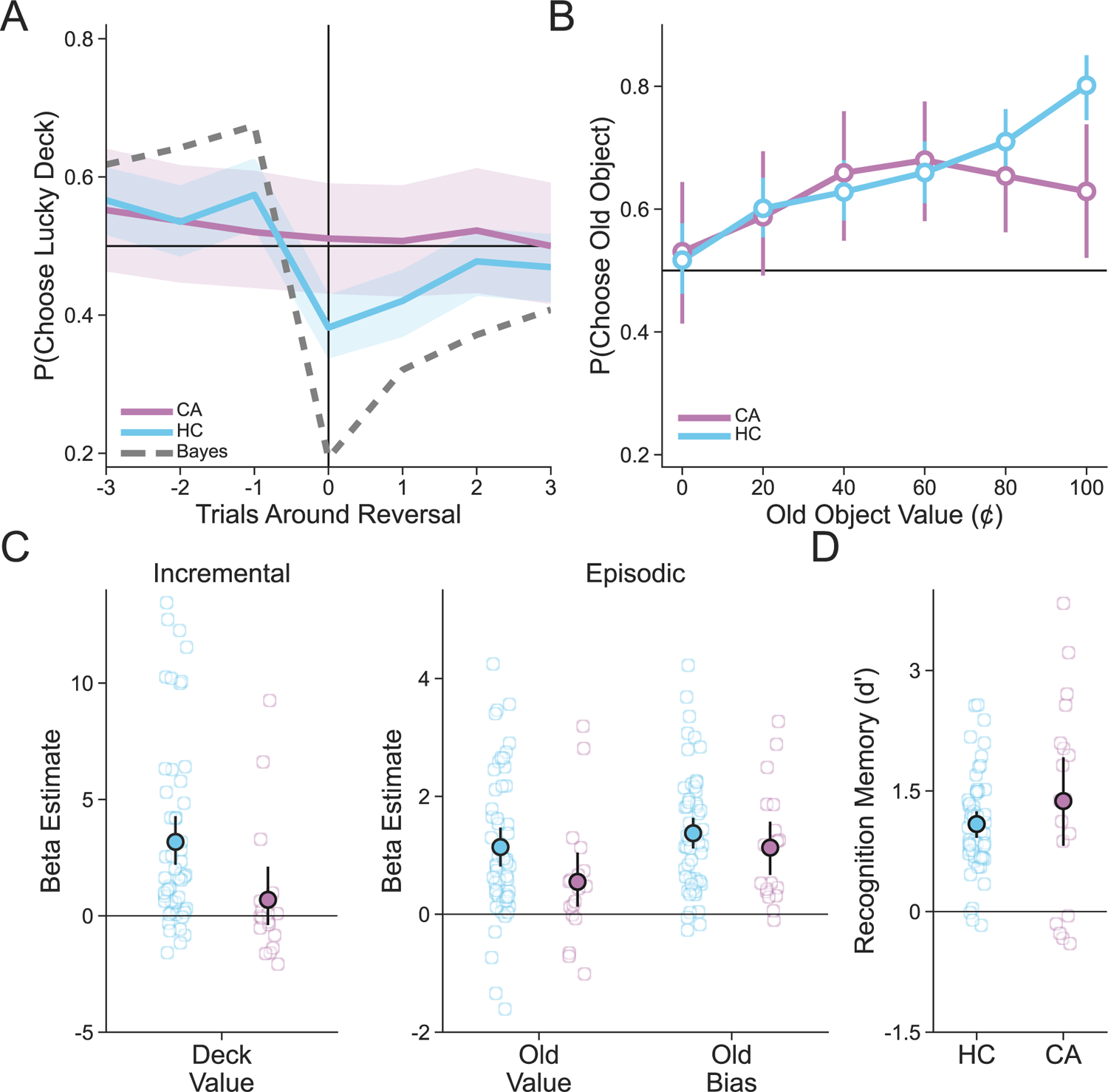
Performance on the multiple learning strategies task. **(A)** Deck learning performance on the multiple learning strategies task as indicated by the proportion of trials on which the currently lucky deck was chosen as a function of how distant those trials were from a reversal in deck value. Performance for both healthy controls (HC) and CA participants is shown alongside a Bayesian observer with perfect episodic memory for visual comparison. Lines represent group averages and bands represent 95% confidence intervals. **(B)** Object value usage on trials in which a previously seen object appeared. Points represent group averages and error bars represent 95% confidence intervals. **(C)** Inverse temperature estimates from the Hybrid model. Individual points represent estimates for each subject, group-level averages are shown as filled in points and error bars represent 95% confidence intervals. **(D)** Recognition memory performance on the subsequent memory task. Individual points represent each participant’s dprime score, filled in points are group-level averages and error bars are 95% confidence intervals.

We next assessed the extent to which both incrementally constructed value and episodic value contributed to choice in a combined Hybrid choice model. This model combined a standard Q learning model with three inverse temperature parameters that captured each participants’ sensitivity to estimated deck value, the true value of previously seen objects, and a bias toward choosing previously seen objects regardless of their value (**Figure 3B-C**). This model of hybrid choice outperformed a biased responder, which again served as a baseline, for both CA participants and controls as there was no difference between groups in estimated out-of-sample predictive performance (β_*Group*_ = −1.631, 95% *CI* = [−11.425, 8.444]). Importantly, this indicates that the behavior of both CA participants and controls was well described by the hybrid choice model, which is expected if CA participants are unimpaired at episodic value learning. For each group, we then assessed whether sensitivity differed from zero and, if so, concluded that participants in that group made choices that were affected by each possible predictor. While healthy controls incorporated deck value into their decisions (β_34_ = 3.173, 95% *CI* = [2.181, 4.189]), CA participants generally did not (β_45_ = 0.681, 95% *CI* = [−0.668, 2.066]). This reward learning deficit was specific to value acquired incrementally, however, because CA participants and controls were both sensitive to episodic value (β_34_ = 1.373, 95% *CI* = [1.095, 1.654]; β_45_ = 1.13, 95% *CI* = [0.59, 1.643]) and were both similarly biased by previously seen objects regardless of their value (β_34_ = 1.142, 95% *CI* = [0.798, 1.477]; β_45_ = 0.551, 95% *CI* = [0.028, 1.056]). Furthermore, while there were no differences between groups for the effects of either episodic value (β_*Group*_ = 0.244, 95% *CI* = [−0.311, 0.820]) or bias (β_*Group*_ = 0.586, 95% *CI* = [−0.051, 1.246]), healthy controls were indeed more sensitive to learned deck value than CA participants (β_*Group*_ = 2.47, 95% *CI* = [0.362, 4.572]).

We additionally had each participant complete a subsequent memory test for a subset of objects shown during the multiple learning strategies task. There was no difference in recognition memory performance between groups (β_*Group*_ = −0.487, 95% *CI* = [−1.144, 0.157]). This result provides further evidence that CA participants were unimpaired at using episodic memory throughout the task relative to their stark impairments in incremental learning. Lastly, CA participants and healthy controls demonstrated no differences in reaction time on this task (β_*Group*_ = −86.40, 95% *CI* = [−222.28, 44.76]), suggesting that the behavioral differences reported here cannot be attributed to motor slowing in CA.

### Controlling for effects of non-motor deficits and disease subtype

We next sought to ensure that the differences in the tasks reported here were specific to deficits in reward learning rather than general cognitive impairment. Controlling for cognitive impairment is particularly important because recent work ^33, 34^ has suggested that incremental learning experiments tax higher level functions, like executive control and working memory, in addition to learning from reward prediction error. To address this issue and assess possible cognitive impairment, we conducted a battery of neuropsychological measures on CA participants (see **Methods**). Of these, a subset of measures were also completed by healthy controls (**Figure 4**). We found no differences in performance between groups on all measures except for the backwards digit span task, which indexes working memory ability, and on which healthy controls scored higher than CA participants (**Table 2**; β_*Group*_ = −2.57, 95% *CI* = [−4.18, −0.92]). Backwards digit span scores were thus included as covariates in regression analyses where possible (see **Methods**) in order to control for impacts of this performance difference on impairments in incremental learning. To further ensure that CA participants’ deficient incremental learning was not due to broad cognitive impairment, we also repeated all analyses excluding seven CA participants (and their matched controls) with mild cognitive impairment (MCI), as indicated by scoring lower than 26 on the MoCA (**Table 1**). While CA participants with MCI consisted of some of the lowest performing participants in our sample (**Figure 2 —Figure supplement 1, Figure 2—Figure supplement 2**), there we no differences in the results across both tasks when they were excluded. It is therefore unlikely that CA participants’ impaired reward learning ability is due to either working memory deficits or cognitive decline more broadly. A full report of these analyses can be found in **Appendix A**.

**Figure 4.**
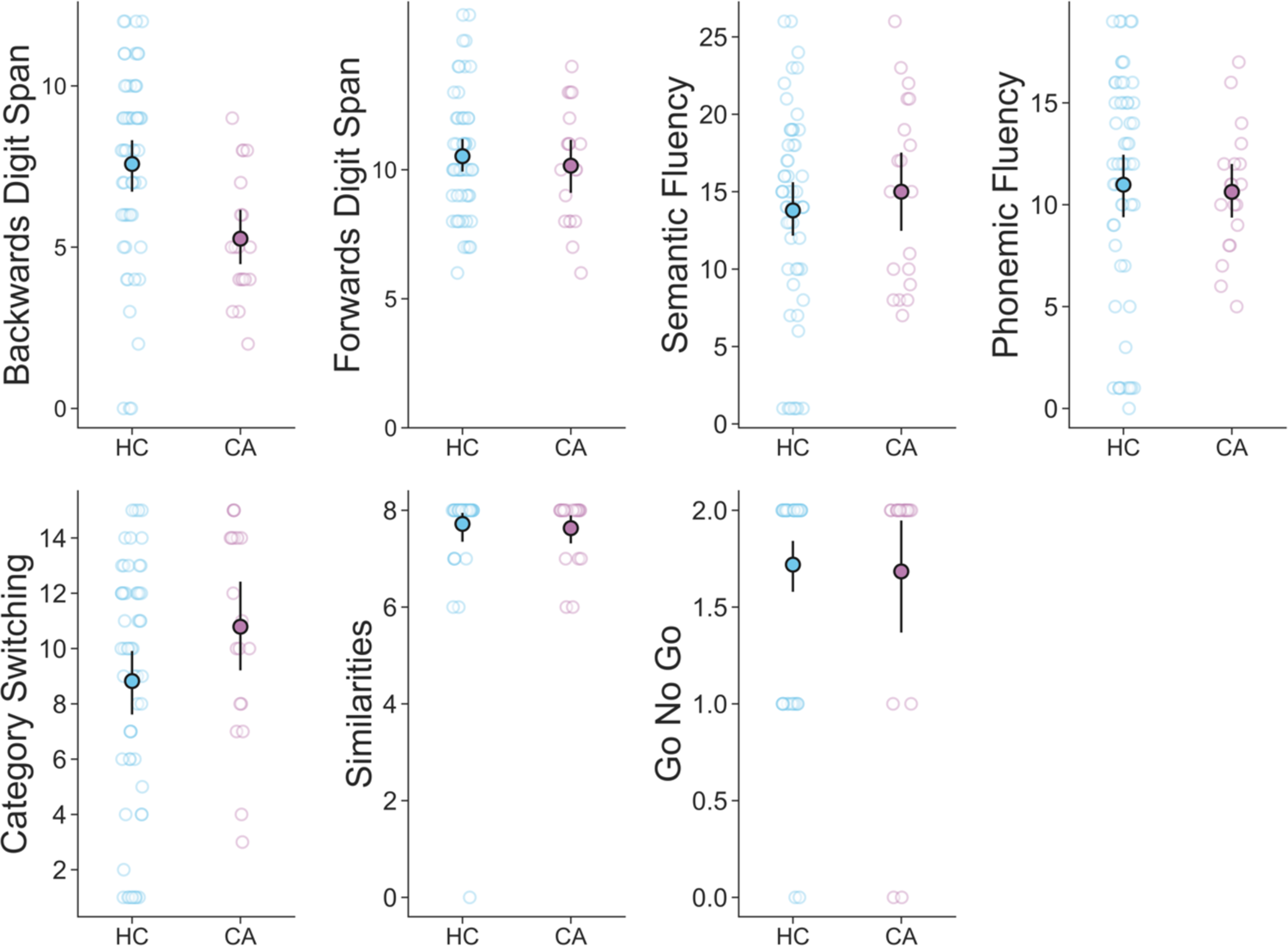
Neuropsychological test performance for the subset of neuropsychological measures that were completed by both CA participants and healthy controls (HC). Scores from individual participants are plotted as empty circles behind group-level means plotted as filled circles with uncertainty represented by 95% confidence intervals.

**Table 1.** Basic CA participant demographics and neuropsychiatric measures

| Participant | Age (years) | Sex | Diagnosis | CCAS | MoCA | BDI | QUIP | FDS | BDS | TMTA (sec) | TMTB (sec) |
| --- | --- | --- | --- | --- | --- | --- | --- | --- | --- | --- | --- |
| Participant 1 | 40 | M | SCA3 | 59 | 21 | 15 | 40 | 8 | 5 | 50 | 150 |
| Participant 2 | 33 | M | SCA3 | 77 | 21 | 9 | 12 | 11 | 5 | 21 | 261 |
| Participant 3 | 20 | F | SCA2 | 92 | 27 | 23 | 10 | 13 | 4 | 41 | 71 |
| Participant 4 | 52 | F | MSA-C | 85 | 27 | 4 | 0 | 8 | 3 | 23 | 158 |
| Participant 5 | 61 | F | SCA2 | 92 | 23 | 15 | 38 | 8 | 4 | 49 | 169 |
| Participant 6 | 56 | M | MSA-C | 100 | 26 | 7 | 8 | 13 | 4 | 66 | 127 |
| Participant 7 | 52 | M | SCA2 | 62 | 26 | 13 | 55 | 11 | 4 | 105 | 287 |
| Participant 8 | 41 | F | SCA2 | 95 | 29 | 18 | 2 | 10 | 8 | 45 | 97 |
| Participant 9 | 43 | M | SCA1 | 87 | 27 | 0 | 6 | 10 | 6 | 52 | 104 |
| Participant 10 | 62 | F | MSA-C | 70 | 21 | 4 | 25 | 6 | 6 | 45 | 121 |
| Participant 11 | 54 | M | SCA2 | 72 | 21 | 0 | 5 | 11 | 5 | 32 | 100 |
| Participant 12 | 67 | F | ILOCA | 101 | 29 | 12 | 16 | 12 | 5 | 51 | 88 |
| Participant 13 | 60 | F | SCA3 | 104 | 28 | 1 | 0 | 14 | 9 | 42 | 113 |
| Participant 14 | 51 | F | SCA10 | 74 | 25 | 13 | 8 | 8 | 2 | 35 | 83 |
| Participant 15 | 66 | M | SCA1 | 60 | 23 | 4 | 20 | 7 | 3 | 66 | 127 |
| Participant 16 | 49 | M | IMCA | 86 | 28 | 17 | 6 | 13 | 7 | 81 | 225 |
| Participant 17 | 54 | M | FA | 98 | 26 | 24 | 8 | 10 | 8 | 50 | 80 |
| Participant 18 | 33 | F | FA | 84 | 27 | 3 | 26 | 9 | 4 | 38 | 84 |
| Participant 19 | 54 | F | IMCA | 113 | 28 | 18 | 4 | 11 | 8 | 33 | 62 |

SCA – Spinocerebellar ataxias, MSA-C – Multiple system atrophy, cerebellar type, ILOCA – Idiopathic late onset cerebellar ataxia, IMCA – Immune-mediated cerebellar ataxia, FA – Friedreich’s ataxia, CCAS – Cerebellar cognitive affective/Schmahmann syndrome scale, MoCA – Montreal cognitive assessment, BDI – Beck’s depression inventory, QUIP – Questionnaire for impulsive-compulsive disorders in Parkinson’s disease, FDS – Forward digit span, BDS – Backward digit span, TMTA – Trail making test part A, TMTB – Trail making test part B

**Table 2.** Neuropsychological test regression analysis results

| Measure | $\beta$ Estimate | 95% Credible Interval |
| --- | --- | --- |
| **Backwards** | **-2.57** | **[-4.18, -0.92]** |
| Forwards | -0.09 | [-1.35, 1.15] |
| Semantic | 1.54 | [-2.12, 5.14] |
| Phonemic | 0.04 | [-2.75, 2.88] |
| Category | 1.68 | [-0.76, 4.10] |
| Similarities | -0.20 | [-0.50, 0.12] |
| Go No Go | 0.0001 | [-0.33, 0.33] |

Results of regression analyses assessing differences in neuropsychological test performance between CA participants and healthy controls.

We next sought to further characterize the nature of CA participants’ reward learning impairment by looking at the relationship between incremental learning sensitivity, as measured by the Q learning models in each task, and performance on our neuropsychological battery. The extent to which CA participants learned about cues in the incremental learning task related only to total CCAS score (*r* = 0.84, *p* < 0.001, *Bonferroni corrected***Table 3**), suggesting that the specific contributions of the cerebellum to cognition may impact performance in this task. The CCAS scale was recently developed to measure the exact types of cognitive impairment that result from damage to the cerebellum^35^. Because more focal cerebellar lesions tend to lead to lower total CCAS scores^36^, this provides further evidence of the necessity for the cerebellum to successfully perform the incremental learning task. The relationship between total CCAS score and performance was driven by the timed portions of the CCAS scale (e.g. the Semantic Fluency and Category Switching measures; **Table 3—Table supplement 1**), suggesting a potential effect of slowed responses in the incremental learning task. While CA participants did indeed respond more slowly than healthy controls on this task (see above), we controlled for this difference in our behavioral analysis. Finally, there was no relationship between any measure and incremental learning ability in the multiple learning strategies task (**Table 3**).

**Table 3.** Correlations between CA participant-level measures and incremental value sensitivity

| Measure | Pearson's R | P Value (Bonferroni Corrected) | Task |
| --- | --- | --- | --- |
| Symptom Duration | -0.0661 | 1 | Incremental Learning |
| SARA (Total) | 0.1067 | 1 | Incremental Learning |
| MoCA (Total) | 0.5225 | 0.1885 | Incremental Learning |
| **CCAS (Total)** | **0.8419** | **0.0001\*** | **Incremental Learning** |
| BDI (Total) | 0.2977 | 1 | Incremental Learning |
| QUIP (Total) | -0.3892 | 0.7356 | Incremental Learning |
| Symptom Duration | -0.3699 | 0.7848 | Multiple Learning Strategies |
| SARA (Total) | -0.2141 | 1 | Multiple Learning Strategies |
| MOCA (Total) | 0.1438 | 1 | Multiple Learning Strategies |
| CCAS (Total) | 0.3849 | 0.6887 | Multiple Learning Strategies |
| BDI (Total) | 0.4641 | 0.3141 | Multiple Learning Strategies |
| QUIP (Total) | -0.074 | 1 | Multiple Learning Strategies |

CA participant-level measures consist of total neuropsychological scores and symptom duration and incremental value sensitivity consists of estimates by the Q Learning model in the incremental learning task and as by the Hybrid Q Learning model in the multiple learning strategies task. CCAS – Cerebellar cognitive affective/Schmahmann syndrome scale, SARA – Scale for the Assessment and Rating of Ataxia, MoCA – Montreal cognitive assessment, BDI – Beck's depression inventory, QUIP – Questionnaire for impulsive-compulsive disorders in Parkinson's disease. \*\*\*p < 0.001

Finally, we addressed the possibility that the subset of our sample of CA participants consisting of diagnoses that were less restricted to the cerebellum, namely the three individuals with multiple system atrophy (MSA) and the two individuals with Friedrich’s ataxia (FA), could be responsible for the deficits reported here. We repeated all analyses with these five participants excluded and found no differences in the results (**Figure 2—Figure supplement 1, Figure 2—Figure supplement 2**). A full report of these analyses can be found in **Appendix B**.

## Discussion

The results of the present work demonstrate that individuals with cerebellar dysfunction, represented by CA cases in our cohort, are impaired at reward learning. While the cerebellum and basal ganglia have traditionally been treated as making separate contributions to learning^5, 21^, recent findings have called this dichotomy into question^8–19^. This work has suggested that, alongside its role in motor learning, the cerebellum likely operates in concert with the basal ganglia to support reinforcement learning from reward. Our study corroborates these findings from animal models^10–, 19^, providing evidence that the human cerebellum is necessary for learning associations from reward. In comparison to age- and sex-matched healthy controls, CA participants were impaired at reward-based learning from trial-and-error. Further, CA participants retained the ability to employ an alternative strategy based in episodic memory to guide their decisions, demonstrating that this impairment is specific to incremental learning. These results challenge the idea that the cerebellum is used primarily for motor learning and shed light on how multiple neural systems may interact with one another to support learning in the non-motor domain.

Our findings join a litany of recent research suggesting that the cerebellum plays a broad role in human cognition^22, 37–39^. Indeed, individuals with damage to the cerebellum demonstrate impairment in a wide range of cognitive functions including cognitive control^40^ and impulsivity^41^. Human functional neuroimaging studies have also revealed cerebellar activity in a variety of different non-motor tasks^22, 38^. Many of these functions are likely supported by the robust bidirectional connections the cerebellum shares with the prefrontal cortex^42, 43^. In particular, recent findings have indicated that individuals with CA have heightened domain-specific impulsive and compulsive behaviors, which is a common symptom of underlying reward system dysfunction^44, 45^. Our study adds to this work by suggesting that the cerebellum is additionally necessary for reward learning in humans.

While there is growing evidence validating the implication of the cerebellum in reward-based learning in animals, there is only limited work on this topic in humans. Early imaging studies, for example, demonstrated cerebellar BOLD activity in patients with substance use disorder who performed reward-based learning tasks^23^ and experienced cravings^24^, and also in response to unexpected reward^25^. However, it remains unknown how cerebellar damage impacts reward learning, as investigations of reward learning in the cerebellum are rare. While two previous studies employed reward-based experimental tasks in individuals with isolated ischemic lesions of the cerebellum^46, 47^, results until this point have remained far from conclusive. Thoma et al. (2008) used a reward-based learning task consisting of an initial acquisition phase in which eight participants with cerebellar damage were rewarded for learning associations between colors and symbols followed by a reversal portion in which they had to disremember previously acquired knowledge and learn new associations for each cue. While participants with cerebellar damage demonstrated no impairment at acquiring new, reward-based knowledge, they were selectively impaired at learning from a single reversal. While this study complements our findings, we found evidence for more global impairment: CA participants in both of our tasks were unable to learn associations from reward on a trial-by-trial basis. Rustemeier et al. (2016) took a different approach by asking twelve individuals with cerebellar damage to learn a simple acquisition task from probabilistic feedback and subsequently transfer this knowledge to re-arranged stimuli. While participants were unimpaired behaviorally at this task, electroencephalographic (EEG) results revealed that they may process reward-based feedback differently from controls. Our findings support this interpretation and further suggest that processing of trial-by-trial feedback is not just different, but impaired, in individuals with cerebellar damage. Finally, while other related studies showed impairment in learning from reinforcement in participants with cerebellar damage^20, 48^, this work has focused primarily on movement-dependent deficiencies.

While our findings suggest that the cerebellum is necessary for incremental reward learning, they cannot speak to the neural circuitry underlying this role. One intriguing possibility is that the cerebellum may operate in tandem with the basal ganglia—canonically seen as the seat of reinforcement learning in the brain^5, 21^—to learn about reward incrementally. Reward prediction error signals in midbrain dopamine neurons that provide input to the basal ganglia^27, 29^ have also been found to be encoded by cerebellar neurons^9, 15, 17, 19^. Further, through excitatory projections to the ventral tegmental area, the cerebellum has widespread reciprocal connections with the basal ganglia and has recently been shown to influence reward-driven behavior through these projections^8, 49^. While reinforcement learning via the basal ganglia and supervised learning via the cerebellum have typically been treated as fulfilling entirely separate roles^5, 21^, these systems appear to be more interdependent than previously thought. Future investigations of the relationship between the basal ganglia and cerebellum are needed to clarify the exact mechanisms underlying reinforcement learning in the brain.

Lastly, there are several potential limitations related to the nature of our sample that should be considered when interpreting these findings. First, cerebellar dysfunction in our sample of CA participants was caused by several different conditions. While most of these pathologies are predominantly restricted to the cerebellum, non-cerebellar brain areas and circuits could also be affected, particularly in participants diagnosed with either MSA or FA. There was, however, no change in the reported reward-based learning deficits when these participants were excluded. Second, while cognitive impairment due to neurodegenerative disease could potentially contribute to some of the deficits measured here, we accounted for this possibility by establishing that the incremental reward learning deficits reported here persist regardless of MCI status. We also collected basic neuropsychological measures from all participants, and CA participants were not different from controls on the vast majority of measures. We focused particularly on possible contributions of working memory given recent work suggesting that working memory plays an important role in incremental reward learning^33, 34^. While CA participants and controls performed similarly on the forward digit span task, CA participants were somewhat impaired at backwards digit span. We controlled for this difference by including backwards digit span scores as covariates in our analyses. Finally, while our control participants completed the study online, we accounted for potential variability caused by this difference in setting by collecting three matched controls for each CA participant in our sample.

Taken together, our findings suggest that the human cerebellum is necessary for reward learning. These results provide new constraints on models of non-motor learning and suggest that the cerebellum and basal ganglia work in tandem to support learning from reinforcement.

## Materials and Methods

### Cerebellar Ataxia Participants

Nineteen individuals with cerebellar ataxia were recruited from the Ataxia Clinic, Columbia University Medical Center and completed both tasks (see **Table 1** for information about basic CA participant demographics and diagnoses). Due to hardware issues, data from one participant on each task was not saved. The first CA participant also completed a shorter pilot version of the incremental learning task, and several changes were made before running this task on the other 18 CA participants. Thus, the final sample for the incremental learning task was 17 CA participants, and the final sample for the multiple learning strategies task was 18 CA participants. Task order was counterbalanced. A neuropsychological battery comprised of the Montreal Cognitive Assessment (MOCA), Beck’s Depression Inventory (BDI), MESA digit forward and backward span, trail making test A and B, and the cerebellar cognitive affective syndrome scale (CCAS) was conducted between tasks for each participant. This battery was specifically selected based on the current understanding of the cerebellum’s role and association with non-motor symptoms, such as depression^50^, executive function^51, 52^, and attention^53^.

### Healthy Controls

Age- and sex-matched participants were recruited through Amazon Mechanical Turk using the Cloud Research Approved Participants feature^54^. To account for potential variability due to online data collection, three matched controls were collected for each CA participant, bringing the total number of controls to 57 (3:1 match). Data from one control was excluded for the multiple learning strategies task due to random responding. Task order was counterbalanced such that the tasks were completed in the identical order to each control’s matched CA participant. A modified online neuropsychological battery consisting of 7 measures was completed in between each task for comparison to individuals with CA. Five of these measures (Semantic Fluency, Phonemic Fluency, Category Switching, Similarities and Go No Go) were directly taken from the CCAS, and two others were comprised of the MESA digit forward back backward span. Participant recruitment was restricted to the United States. Before starting each task, all participants were required to score 100% on a quiz that tested their comprehension of the instructions. Informed consent was obtained with approval form the Columbia University Institutional Review Board.

### Experimental Design

#### Incremental Learning Task

In the incremental learning task (**Figure 1A**), participants were told that they would be playing a game where they were required to press a key, either F or J, whenever one of two symbols was seen, and that they would receive feedback about whether they had pressed correctly following each trial. They were then informed that it was their job to determine which key they should press for each symbol, and that what key is best will change throughout the experiment. Outcomes were determined by a drifting probability such that each button was correct for each image 50% of the time. Critically, these probabilities differed over time, thus encouraging constant learning throughout the task. Participants were told to press the F key with their left index finger and the J key with their right index finger. The response period during which the symbol remained on the screen lasted 1.5 seconds, with feedback displayed for 1 second immediately following the response period. An intertrial interval featuring a fixation cross was shown for an average of 1 second, but varied between 0.5 and 1.5 seconds. Lastly, to provide a rewarding outcome for correct responses, participants were informed that they could earn bonus money based on their performance. Correct responses were worth an additional cent each.

#### Multiple Learning Strategies Task

The other task completed by participants was previously developed by our lab ^31, 32^ to measure the relative contribution of incremental learning and episodic memory to decisions (**Figure 1B**). Participants were told that they would be playing a card game where their goal was to win as much money as possible. Each trial consisted of a choice between two decks of cards that differed based on their color (red or blue). Participants had two seconds to decide between the decks. The outcome of each decision was then immediately displayed for 1.5 seconds. Following each decision, participants were shown a fixation cross during the intertrial interval period which varied in length (mean = 1.5 seconds, min = 1 seconds, max = 2 seconds). Decks were equally likely to appear on either side of the screen on each trial. Participants completed a total of 150 trials.

Participants were made aware that there were two ways they could earn bonus money throughout the task, which allowed for the use of incremental learning and episodic memory respectively. First, at any point in the experiment one of the two decks was “lucky”, meaning that the expected value (*V*) of one deck color was higher than the other (*V*_*lucky*_=63¢, *V*_unlucky_=37¢). Outcomes ranged from $0 to $1 in increments of 20¢. Critically, the mapping from *V* to deck color reversed periodically throughout the experiment, which incentivized participants to utilize each deck’s recent reward history to determine the identity of the currently lucky deck. Second, to assess the use of episodic memory throughout the task, each card within a deck featured an image of a trial-unique object that could re-appear once throughout the experiment after initially being chosen. Participants were told that if they encountered a card a second time it would be worth the same amount as when it was first chosen, regardless of whether its deck color was currently lucky or not. On a given trial *t*, cards chosen once from trials *t* − 9 through *t* − 30 had a 60% chance of reappearing following a sampling procedure designed to prevent each deck’s expected value from becoming skewed by choice, minimize the correlation between the expected value of previously seen cards and deck expected value, and ensure that choosing a previously selected card remained close to 50¢.

Following completion of the multiple learning strategies task, we tested participants’ memory for the trial-unique objects. Participants completed up to 54 three-part memory trials. An object was first displayed on the screen and participants were asked whether or not they had previously seen the object and were given five response options: Definitely New, Probably New, Don’t Know, Probably Old, Definitely Old. If the participant indicated that they had not seen the object before or did not know, they moved on to the next trial. If, however, they indicated that they had seen the object before they were then asked if they had chosen the object or not. Lastly, if they responded that they had chosen the object, they were asked what the value of that object was.

### Computational Models

In order to capture subjective estimates of incrementally constructed value on each task, we fit computational models to participants’ choices. Below we describe each of these models in detail.

#### Q Learning Models

We modeled incremental reward learning using a Q Learning model, which is a standard model-free reinforcement learner that assumes a stored value (*Q*) for each deck is updated over time ^26, 28^.

*Q* is then referenced on each decision in order to guide choices. After each outcome, *r*., the value for an option *Q*_1_is updated according to the following rule if that option is chosen:

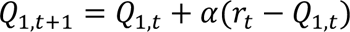

And is not updated if a different option is chosen:

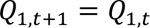

Likewise, if a different option is chosen, its value is updated equivalently. Large differences between estimated value and outcomes therefore have a larger impact on updates, but the overall degree of updating is controlled by the learning rate, α, which is a free parameter constrained to lie between 0 and 1.

For the incremental learning task, the model learned separate Q values for each cue and button combination, such that four Q values were estimated in total. Decisions were then modeled using the following rule:

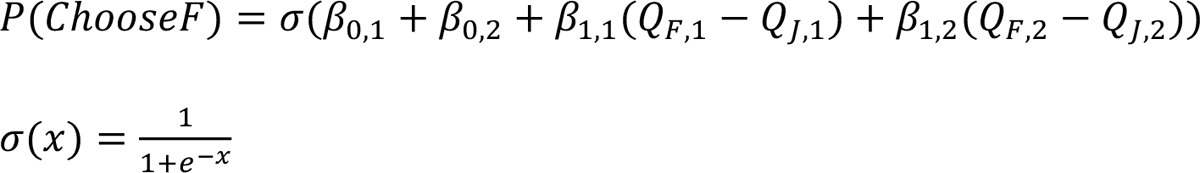

such that four inverse temperatures β were estimated to capture a bias toward choosing a key for each cue (β_1,-_and β_1,<_) and sensitivity to incrementally learned value for each cue (β_-,-_ and β_-,<_). This model is referred to as the “Q Learner” model throughout the text.

For the multiple learning strategies task, the model learned separate Q values for each deck color, such that two Q values were estimated in total. Decisions were then modeled using the following rule:

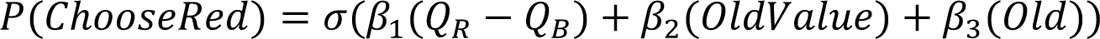

such that three inverse temperatures β were estimated to capture sensitivity to incrementally learned value (β_-_), sensitivity to the value of previously seen objects (β_1_), and a bias toward choosing the deck featuring a previously seen object regardless of its value (β_2_). The predictor *OldValue* was the coded true value of a previously seen object (ranging from 0.5 if the value was $1 on the red deck or $0 on the blue deck to −0.5 if the value was $0 on the red deck and $1 on the blue deck) and the predictor *Old* was coded as 0.5 if the red deck featured a previously seen object and −0.5 if the blue deck did instead. For both of these predictors, trials that did not feature a previously seen object were coded as 0. This model is referred to as the “Hybrid” model throughout the text.

#### Biased Responder Model

For both tasks, we compared the performance of the Q Learning models to a model which made choices that were completely independent of reward information. For the incremental learning task, this model was simply:

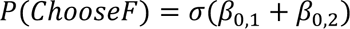

such that choices depended only on choosing a button to press for each cue throughout the experiment. For the multiple learning strategies task, this model was:

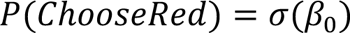

such that choices depended only on preferring one deck over the other throughout the experiment. Our logic in using this model as a baseline was that responses captured by the Q learning models should, at a minimum, outperform a biased responder that did not consider reward in order for it to make meaningful predictions about participants’ behavior.

#### Posterior Inference and Model Comparison

Model parameters for each participant were estimated using Bayesian inference. The joint posterior was approximated using No-U-Turn Sampling^55^ as implemented in stan^56^. Four chains with 2000 samples (1000 discarded as burn-in) were run for a total of 4000 posterior samples per model per subject. Chain convergence was determined by ensuring that the Gelman-Rubin statistic *R*^S^ was close to 1 for all parameters. For the incremental learning task, the Q learner did not converge for one CA participant, and so that individual and their matched controls were removed from further model-based analyses. For the multiple learning strategies task, all models for all participants converged.

Under this approach, the likelihood function for all models can be written as:

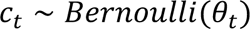

where *c*_t_ is 1 if the subject chose F (in the resonse mapping task) or red (in the multiple learning strategies task). Here, θ_t_ is the linear combination of inverse temperature parameters and predictors explained above for each model. For the Q learning models, the learning rate,α, had the following weakly informative prior:

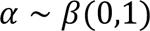

For all models, every inverse temperature parameter had the following weakly informative prior:

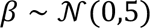

Model fit was assessed using approximate leave-one-out cross validation estimated using Pareto-smoothed importance sampling^57^. The expected log pointwise predictive density (ELPD) was computed and used as a measure of out-of-sample predictive fit for each model.

#### Bayesian Observers

In order to provide a normative performance benchmark, we simulated beliefs about incremental value as estimated by Bayesian observers for each task. For the incremental learning task, this learner was a Kalman Filter^58^ and for the multiple learning strategies task this learner was a reduced Bayesian change-point detection model^59^. Choices in the incremental learning task were made according to which button the observer believed was the most likely to be rewarded for each cue at each time point. Choices in the multiple learning strategies task were made differently depending on whether a previously seen object was present. For trials in which no previously seen object was shown, the observer responded according to its beliefs about deck value. For trials in which a previously seen object was present, however, the observer compared the value of that object to its belief about deck value for the opposing deck and chose accordingly. In this way, the observer was augmented with “perfect” episodic memory.

#### Regression Models

Mixed effects Bayesian regressions were used to test effects of group (CA participant or control). Group membership was allowed to vary randomly by CA participant identifier, *pid*, such that CA participants and matched controls were assigned the same ID. In these models, *GroupID* was coded as −0.5 for CA participants and 0.5 for controls. We additionally controlled for working memory ability by including backwards digit span scores, *dsBwd*, as a standardized covariate in these analyses.

For the incremental learning task, we assessed behavioral performance using the following logistic regression:

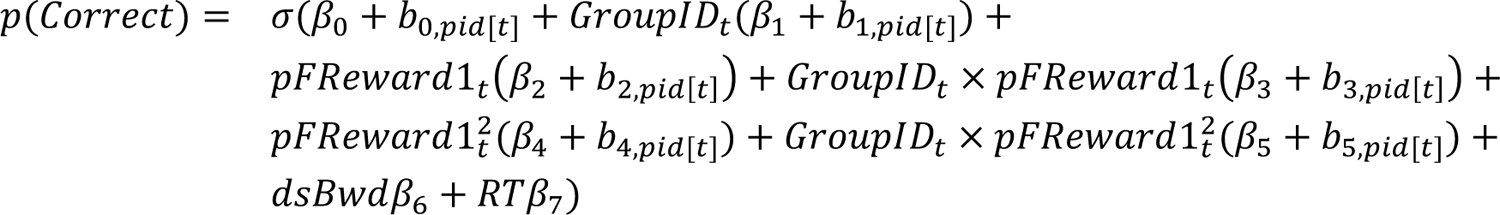

Here, and in all regressions described in this section, β stands for fixed effects and *b* stands for random effects of CA participant ID. The predictor *pFReward1* indicates the true underlying difficulty of the task and is the probability that the F key was rewarding for cue one. A second-order polynomial was included for this predictor as extreme values indicate portions of the task that are easier and middling values indicate portions of the task that were more difficult. Interaction effects of this predictor and group were included to capture differences in sensitivity to the underlying task difficulty between the groups. Lastly, the reaction time, *RT*, on each decision was included as a standardized covariate in this analysis to account for any differences that may be due to slowed responding by individuals with CA on this task.

For both the incremental learning and multiple learning strategies tasks, we assessed whether there were differences between the groups on Q learning model performance compared to the baseline biased responder model with the following linear regression:

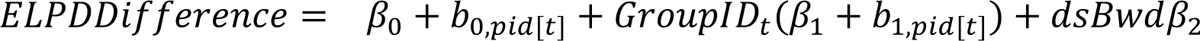

where *ELPDDifference* was the difference in model performance (Q Learning model ELPD - Biased Responder ELPD; see above) for each subject.

For the multiple learning strategies task, we assessed behavioral incremental learning performance using the following logistic regression:

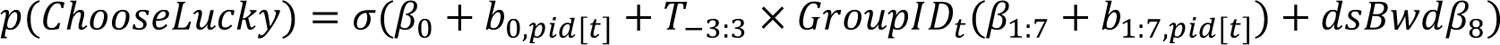

In this regression, we grouped trials according to their distance from a reversal, up to three trials prior to (*t* = −3: −1), during (*t* = 0), and after (*t* = 1: 3) a reversal occurred. We then dummy coded them to measure their effects on the degree to which the lucky deck was chosen and interacted each dummy coded regressor with group to measure how this was affected by group membership.

We then assessed the degree to which each group used either incrementally learned deck value, the value of previously seen objects, or a bias toward previously seen objects regardless of their value as estimated by the Hybrid Q learning model using a simple linear regression of the following form for each of these inverse temperature parameters and groups:

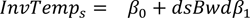

Here we interested primarily in the intercept, β_1_, as this determined the degree to which each group’s inverse temperatures were above zero. We additionally assessed differences between groups on each of these measures by including fixed and random effects for group that varied by matched participant ID, as in previously described regression analyses.

We also assessed the impact of group on subsequent memory performance following the multiple learning strategies task using the following linear regression:

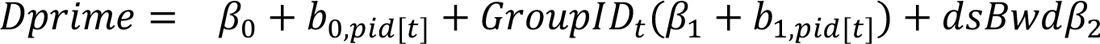

where *Dprime* is the signal detection measure *d*′, which is the difference in z scored hit rate and false alarm rate for each participant.

We were also interested in determining whether there were any differences in reaction times between individuals with CA and matched controls due to motor impairment. For both tasks, we did this by assessing whether there were any differences in reaction time between groups:

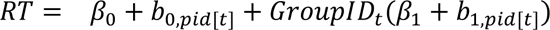

where *RT* was the median reaction time across trials in either task. A separate regression of this form was used for each of the two tasks. We also assessed whether there were differences on each neuropsychological measure using a similar regression.

For all regression analyses, fixed effects are reported in the text as the mean of each parameter’s marginal posterior distribution alongside 95% credible intervals, which indicate where 95% of the posterior density falls. Parameter values outside of this range are unlikely given the model, data, and priors. Thus, if the range of likely values does not include zero, we conclude that a meaningful effect was observed.

### Code Accessibility

All code used to analyze the data in this study may be found here: https://github.com/boomsbloom/ataxia-rl

## Acknowledgements

J.N. was supported by the NSF Graduate Research Fellowship (1644869). S.H.K. was supported by NINDS R01NS104423, NINDS R01 NS118179, NINDS R01 NS124854, and National Ataxia Foundation. D.S. was supported by an NSF CRCNS award (1822619), NIMH R01 MH121093 and the Kavli Foundation.

## Supplementary Information

**Figure 2—Figure supplement 1.**
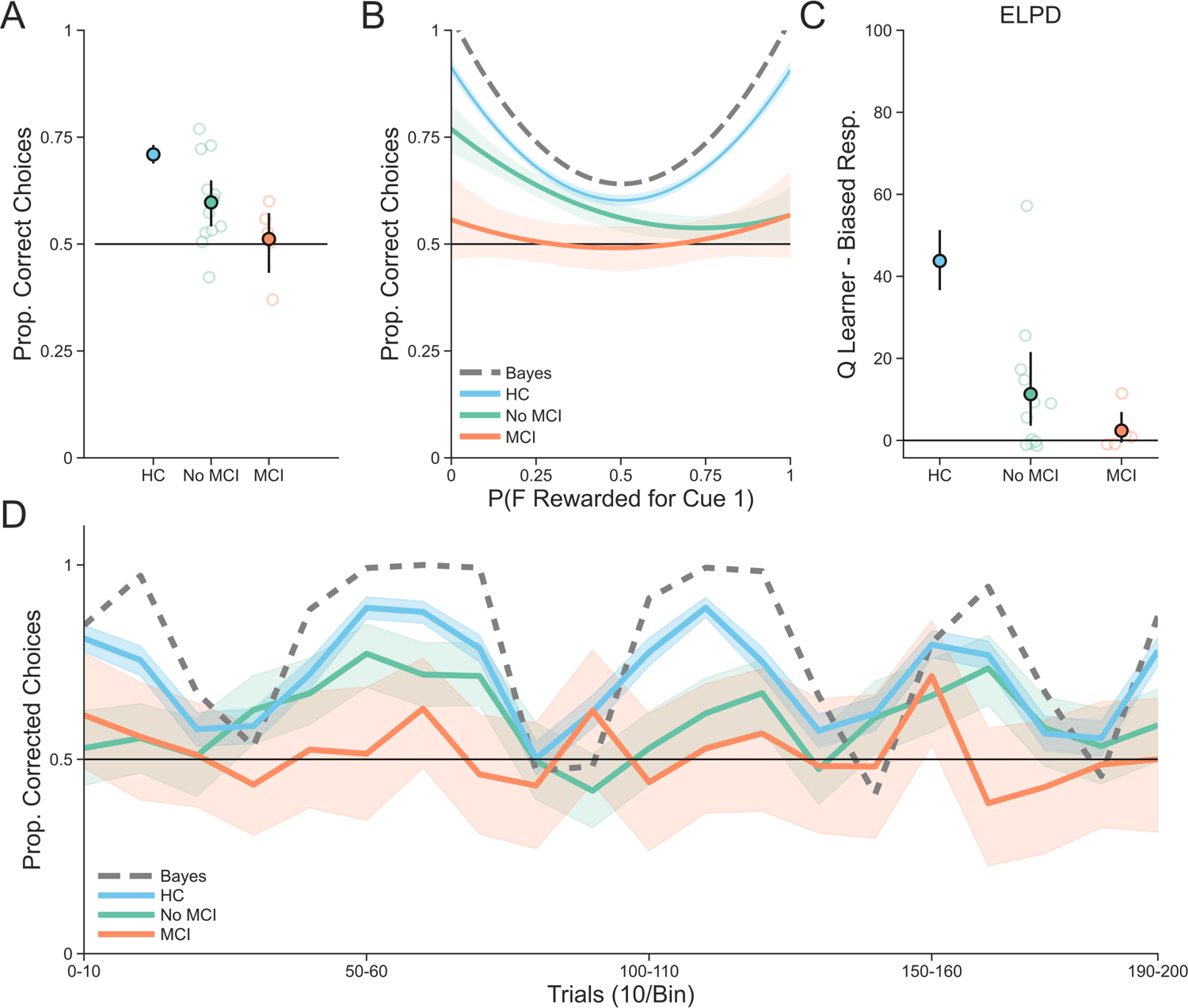
CA participant performance on the incremental learning task separated by mild cognitive impairment (MCI versus all others) and compared to healthy controls (HC). **(A)** Performance on the incremental learning task averaged across all trials. Individual points are averages for each subject and filled in points represent group-level averages. Error bars are 95% confidence intervals. **(B)** Performance on the incremental learning task as a function of task difficulty, which is indexed by the true underlying probability that pressing the F key was the correct response (>50%) on each trial. Points represent group level averages from 13 bins with an equal number of trials, lines represent the fit of a second-order linear model, and error bars and bands represent 95% confidence intervals. **(C)** The difference in estimated out-of-sample predictive performance (as measured by expected log pointwise predictive density; ELPD) between the Q Learner and the Biased Responder model for each group. Individual points are the ELPD difference for each subject and filled in points represent group-level averages. Error bars are 95% confidence intervals. **(D)** Performance on the incremental learning task over time. Each timepoint represents ten trials. Lines are group averages and bands are 95% confidence intervals. For normative comparison, the performance of the Bayesian observer is shown as a dotted gray line.

**Figure 2—Figure supplement 2.**
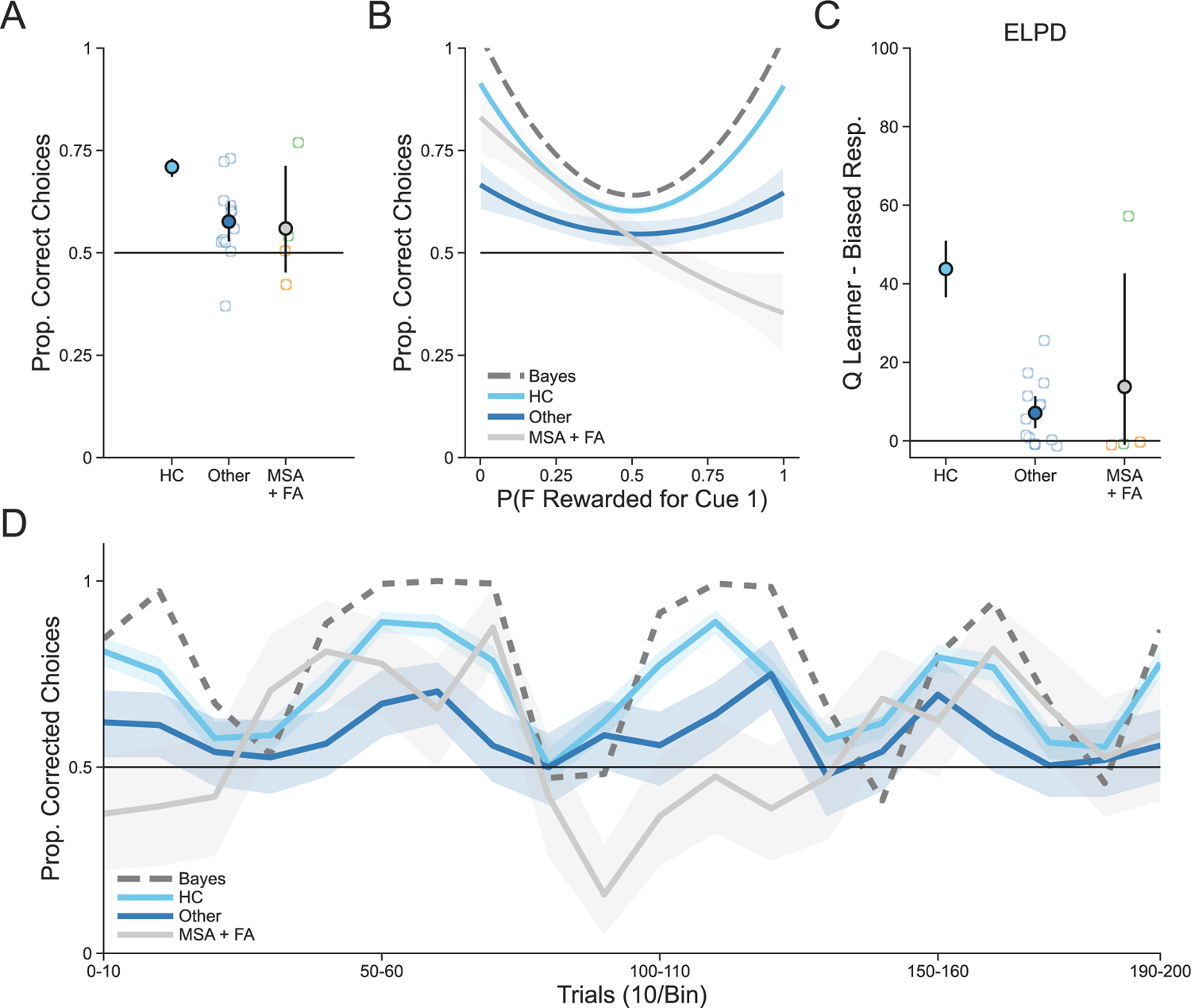
CA participant performance on the incremental learning task separated by diagnosis (MSA + FA individuals versus all others) and compared to healthy controls (HC). **(A)** Performance on the incremental learning task averaged across all trials. Individual points are averages for each subject and filled in points represent group-level averages. Error bars are 95% confidence intervals. MSA participants are shown in green and FA participants are shown in orange. **(B)** Performance on the incremental learning task as a function of task difficulty, which is indexed by the true underlying probability that pressing the F key was the correct response (>50%) on each trial. Points represent group level averages from 13 bins with an equal number of trials, lines represent the fit of a second-order linear model, and error bars and bands represent 95% confidence intervals. **(C)** The difference in estimated out-of-sample predictive performance (as measured by expected log pointwise predictive density; ELPD) between the Q Learner and the Biased Responder model for each group. Individual points are the ELPD difference for each subject and filled in points represent group-level averages. Error bars are 95% confidence intervals. **(D)** Performance on the incremental learning task over time. Each timepoint represents ten trials. Lines are group averages and bands are 95% confidence intervals. For normative comparison, the performance of the Bayesian observer is shown as a dotted gray line.

**Figure 3—Figure supplement 1.**
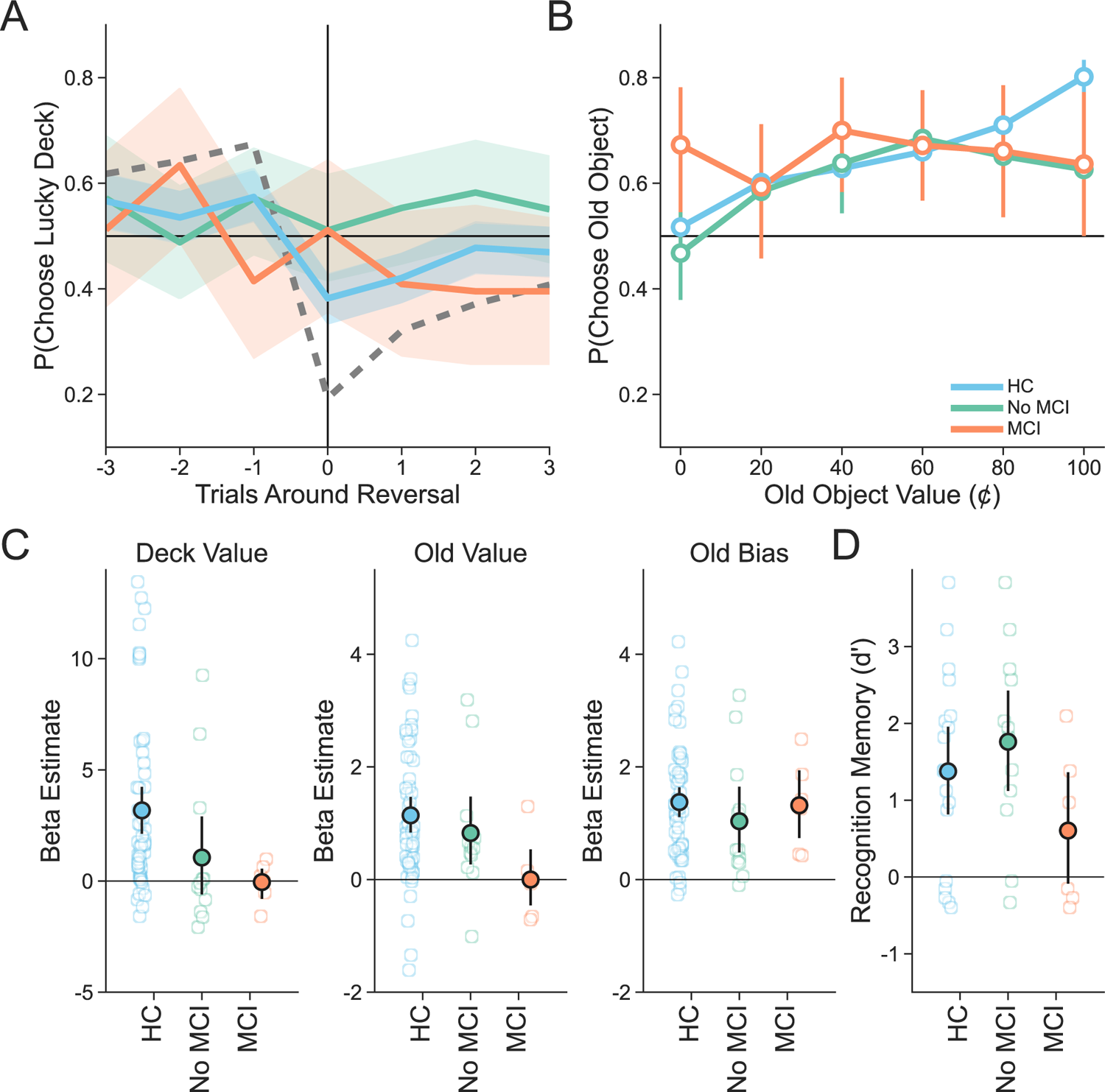
CA participant performance on the multiple learning strategies task separated by mild cognitive impairment (MCI versus all others) and compared to healthy controls (HC). **(A)** Deck learning performance on the multiple learning strategies task as indicated by the proportion of trials on which the currently lucky deck was chosen as a function of how distant those trials were from a reversal in deck value. Lines represent group averages and bands represent 95% confidence intervals. **(B)** Object value usage on trials in which a previously seen object appeared. Points represent group averages and error bars represent 95% confidence intervals. **(C)** Inverse temperature estimates from the Hybrid model. Individual points represent estimates for each subject, group-level averages are shown as filled in points and error bars represent 95% confidence intervals. **(D)** Recognition memory performance on the subsequent memory task. Individual points represent each participant’s dprime score, filled in points are group-level averages and error bars are 95% confidence intervals.

**Figure 3—Figure supplement 2.**
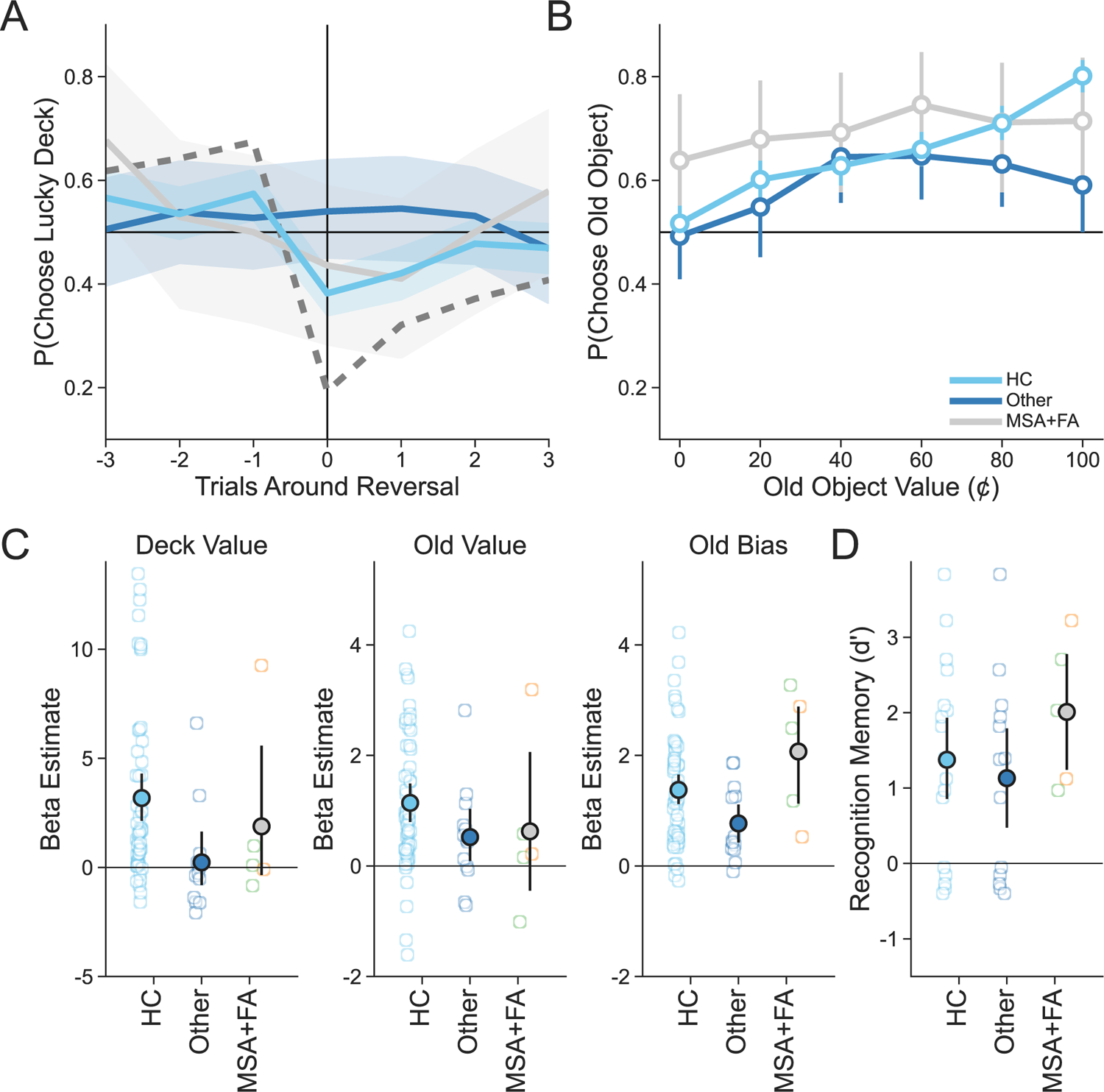
CA participant performance on the multiple learning strategies task separated by diagnosis (MSA + FA individuals versus all others) and compared to healthy controls (HC) **(A)** Deck learning performance on the multiple learning strategies task as indicated by the proportion of trials on which the currently lucky deck was chosen as a function of how distant those trials were from a reversal in deck value. Lines represent group averages and bands represent 95% confidence intervals. MSA participants are shown in green and FA participants are shown in orange. **(B)** Object value usage on trials in which a previously seen object appeared. Points represent group averages and error bars represent 95% confidence intervals. **(C)** Inverse temperature estimates from the Hybrid model. Individual points represent estimates for each subject, group-level averages are shown as filled in points and error bars represent 95% confidence intervals. **(D)** Recognition memory performance on the subsequent memory task. Individual points represent each participant’s dprime score, filled in points are group-level averages and error bars are 95% confidence intervals.

---

**Table 3—Table supplement 1.**
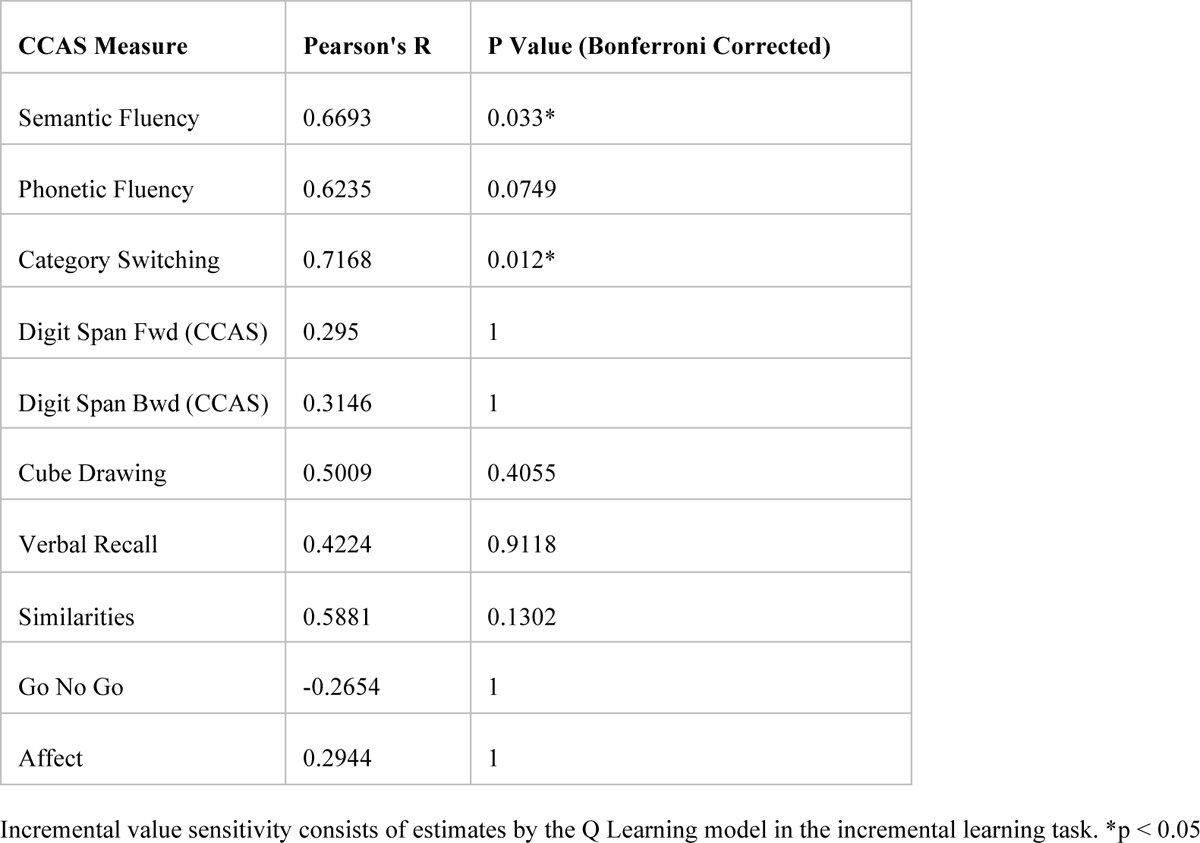
Correlations between CA participant CCAS subscale measures and incremental value sensitivity

| CCAS Measure | Pearson's R | P Value (Bonferroni Corrected) |
| --- | --- | --- |
| Semantic Fluency | 0.6693 | 0.033* |
| Phonetic Fluency | 0.6235 | 0.0749 |
| Category Switching | 0.7168 | 0.012* |
| Digit Span Fwd (CCAS) | 0.295 | 1 |
| Digit Span Bwd (CCAS) | 0.3146 | 1 |
| Cube Drawing | 0.5009 | 0.4055 |
| Verbal Recall | 0.4224 | 0.9118 |
| Similarities | 0.5881 | 0.1302 |
| Go No Go | -0.2654 | 1 |
| Affect | 0.2944 | 1 |

Incremental value sensitivity consists of estimates by the Q Learning model in the incremental learning task. \* $p < 0.05$

## Appendix A

### Results excluding CA participants with MCI

In order to ensure that our results were not biased by patients with mild cognitive impairment (scoring <26 on the MoCA), here we report all primary analyses with these participants (1, 2, 5, 10, 11, 14, and 15) excluded. Further mention of CA participants in this section refers to all CA participants excluding these seven participants and their matched controls.

In the incremental learning task, CA participants made overall fewer correct choices compared to healthy controls (β_*Group*_ = −0.10, 95% *CI* = [−0.02, −0.18]; **Figure 2—Figure Supplement 1A**). CA participants’ choices were also less correct during periods of learning where action-outcome contingencies were more deterministic (e.g. close to 100%) compared to more difficult periods of learning (β_*Group×pFReward1*_^2^ = −5.69, 95% *CI* = [−8.37, −3.20]; **Figure 2—Figure Supplement 1B-C**. Overall, this difference in performance indicates that CA participants did not learn from reward feedback. CA participants responded slightly more slowly than healthy controls on this task (β_*Group*_ = −147.69, 95% *CI* = [−226.03, −72.97]). Controls were also much better fit by the Q learner compared to the biased responder, while this improvement in fit was largely absent for CA participants (β_*Group*_ = 25.54, 95% *CI* = [9.21, 41.42]).

In the multiple learning strategies task, CA participants were less responsive to reward outcomes compared to controls (**Figure 3—Figure Supplement 1A**). Specifically, controls were disrupted more than CA participants by reversals (β_*Group*×./1_ = −0.71, 95% *CI* = [−1.21, −0.22]) and remained below chance performance after a reversal occurred (β_*Group*×./.2-_ = −0.51, 95% *CI* = [−1.00, −0.03]). The model of hybrid choice outperformed the baseline biased responder for both CA participants and controls as there was no difference between groups in estimated out-of-sample predictive performance (β_*Group*_ = −7.01, 95% *CI* = [−20.17, 5.47]). While controls incorporated deck value into their decisions (β_34_ = 3.05, 95% *CI* = [1.87, 4.24]), CA participants generally did not (β_45_ = 1.04, 95% *CI* = [−1.20, 3.22]; **Figure 3—Figure Supplement 1C**). CA participants and controls were both sensitive to episodic value (β_34_ = 1.41, 95% *CI* = [1.69, 1.12]; β_45_ = 1.03, 95% *CI* = [0.31, 1.76]; **Figure 3—Figure Supplement 1B-C**) and were both similarly biased by previously seen objects regardless of their value (β_34_ = 1.18, 95% *CI* = [0.77, 1.58]; β_45_ = 0.83, 95% *CI* = [0.14, 1.53]). CA participants and healthy controls demonstrated no differences in reaction time on this task (β_*Group*_ = −7.91, 95% *CI* = [−173.68, 157.65]). Lastly, there was no difference in recognition memory performance between groups (β_*Group*_ = −0.59, 95% *CI* = [−1.34, 0.157]).

## Appendix B

### Results excluding CA participants with MSA + FA

In order to ensure that our results were not biased by patients with diagnoses that were not restricted to the cerebellum (those with MSA or FA), here was report all primary analyses with these participants (4, 6, 10, 17, 18) excluded. Further mention of CA participants in this section refers to all CA participants excluding these five participants and their matched controls.

In the incremental learning task, CA participants made overall fewer correct choices compared to healthy controls (β_*Group*_ = −1.27, 95% *CI* = [−0.74, −1.82]; **Figure 2—Figure Supplement 2A**). CA participants’ choices were also less correct during periods of learning where action-outcome contingencies were more deterministic (e.g. close to 100%) compared to more difficult periods of learning (β_*Group×pFReward1*_^2^ = −4.76, 95% *CI* = [−6.32, −3.08]; **Figure 2—Figure Supplement 2B-C**. Overall, this difference in performance indicates that CA participants did not learn from reward feedback. CA participants responded slightly more slowly than healthy controls on this task (β_*Group*_ = −112.40, 95% *CI* = [−217.35, −5.03]). Controls were also much better fit by the Q learner compared to the biased responder, while this improvement in fit was largely absent for CA participants (β_*Group*_ = 37.30, 95% *CI* = [20.41, 54.63]).

In the multiple learning strategies task, CA participants were less responsive to reward outcomes compared to controls (**Figure 3—Figure Supplement 2A**). Specifically, controls were disrupted more than CA participants by reversals (β_*Group*×./1_ = −1.15, 95% *CI* = [−1.66, −0.65]) and remained below chance performance after a reversal occurred (β_*Group*×./.2-_ = −0.66, 95% *CI* = [−1.12, −0.20]). The model of hybrid choice outperformed the baseline biased responder for both CA participants and controls as there was no difference between groups in estimated out-of-sample predictive performance (β_*Group*_ = 3.96, 95% *CI* = [−7.29, 14.89]). While controls incorporated deck value into their decisions (β_34_ = 2.70, 95% *CI* = [1.57, 3.92]), CA participants generally did not (β_45_ = 0.22, 95% *CI* = [−1.36, 1.71]; **Figure 3—Figure Supplement 2C**). CA participants and controls were both sensitive to episodic value (β_34_ = 1.31, 95% *CI* = [1.63, 0.98]; β_45_ = 0.77, 95% *CI* = [0.37, 1.15]; **Figure 3—Figure Supplement 2B-C**). Controls were biased by previously seen objects regardless of their value (β_34_ = 1.35, 95% *CI* = [1.75, 0.95]), but CA participants demonstrated this effect less strongly (β_45_ = 0.51, 95% *CI* = [1.09, −0.05]).

CA participants and healthy controls demonstrated no differences in reaction time on this task (β_*Group*_ = −39.00, 95% *CI* = [−192.12, 113.59]). Lastly, there was no difference in recognition memory performance between groups (β_*Group*_ = −0.27, 95% *CI* = [−0.97, 0.470]).

## References

1. Raymond, J. L., Lisberger, S. G. & Mauk, M. D. The Cerebellum: A Neuronal Learning Machine? Science 272, 1126–1131 (1996).

2. Llinás, R. & Welsh, J. P. On the cerebellum and motor learning. Curr. Opin. Neurobiol. 3, 958–965 (1993).

3. Ito, M. & Itō, M. The Cerebellum and Neural Control. (Raven Press, 1984).

4. Marr, D. A theory of cerebellar cortex. J. Physiol. 202, 437–470 (1969).

5. Doya, K. What are the computations of the cerebellum, the basal ganglia and the cerebral cortex? Neural Netw. 12, 961–974 (1999).

6. Wolpert, D. M., Miall, R. C. & Kawato, M. Internal models in the cerebellum. Trends Cogn. Sci. 2, 338–347 (1998).

7. Raymond, J. L. & Medina, J. F. Computational Principles of Supervised Learning in the Cerebellum. Annu. Rev. Neurosci. 41, 233–253 (2018).

8. Caligiore, D., Arbib, M. A., Miall, R. C. & Baldassarre, G. The super-learning hypothesis: Integrating learning processes across cortex, cerebellum and basal ganglia. Neurosci. Biobehav. Rev. 100, 19–34 (2019).

9. Hull, C. Prediction signals in the cerebellum: Beyond supervised motor learning. eLife 9, e54073 (2020).

10. Sendhilnathan, N. & Goldberg, M. E. The mid-lateral cerebellum is necessary for reinforcement learning. http://biorxiv.org/lookup/doi/10.1101/2020.03.20.000190 (2020) doi:10.1101/2020.03.20.000190.

11. Sendhilnathan, N., Semework, M., Goldberg, M. E. & Ipata, A. E. Neural Correlates of Reinforcement Learning in Mid-lateral Cerebellum. Neuron 106, 188–198.e5 (2020).

12. Sendhilnathan, N., Ipata, A. & Goldberg, M. E. Mid-lateral cerebellar complex spikes encode multiple independent reward-related signals during reinforcement learning. Nat. Commun. 12, 6475 (2021).

13. Larry, N., Yarkoni, M., Lixenberg, A. & Joshua, M. Cerebellar climbing fibers encode expected reward size. eLife 8, e46870 (2019).

14. Carta, I., Chen, C. H., Schott, A. L., Dorizan, S. & Khodakhah, K. Cerebellar modulation of the reward circuitry and social behavior. Science 363, eaav0581 (2019).

15. Heffley, W. & Hull, C. Classical conditioning drives learned reward prediction signals in climbing fibers across the lateral cerebellum. eLife 8, e46764 (2019).

16. Wagner, M. J., Kim, T. H., Savall, J., Schnitzer, M. J. & Luo, L. Cerebellar granule cells encode the expectation of reward. Nature 544, 96–100 (2017).

17. Heffley, W. et al. Coordinated cerebellar climbing fiber activity signals learned sensorimotor predictions. Nat. Neurosci. 21, 1431–1441 (2018).

18. Kostadinov, D., Beau, M., Blanco-Pozo, M. & Häusser, M. Predictive and reactive reward signals conveyed by climbing fiber inputs to cerebellar Purkinje cells. Nat. Neurosci. 22, 950–962 (2019).

19. Ohmae, S. & Medina, J. F. Climbing fibers encode a temporal-difference prediction error during cerebellar learning in mice. Nat. Neurosci. 18, 1798–1803 (2015).

20. Therrien, A. S., Wolpert, D. M. & Bastian, A. J. Effective reinforcement learning following cerebellar damage requires a balance between exploration and motor noise. Brain J. Neurol. 139, 101–114 (2016).

21. Doya, K. Complementary roles of basal ganglia and cerebellum in learning and motor control. Curr. Opin. Neurobiol. 10, 732–739 (2000).

22. King, M., Hernandez-Castillo, C. R., Poldrack, R. A., Ivry, R. B. & Diedrichsen, J. Functional boundaries in the human cerebellum revealed by a multi-domain task battery. Nat. Neurosci. 22, 1371–1378 (2019).

23. Volkow, N. D. et al. Expectation enhances the regional brain metabolic and the reinforcing effects of stimulants in cocaine abusers. J. Neurosci. Off. J. Soc. Neurosci. 23, 11461– 11468 (2003).

24. Grant, S. et al. Activation of memory circuits during cue-elicited cocaine craving. Proc. Natl. Acad. Sci. U. S. A. 93, 12040–12045 (1996).

25. Ramnani, N., Elliott, R., Athwal, B. S. & Passingham, R. E. Prediction error for free monetary reward in the human prefrontal cortex. NeuroImage 23, 777–786 (2004).

26. Sutton, R. S. & Barto, A. G. Reinforcement Learning: An Introduction. 352.

27. Houk, J. C., Adams, J. L. & Barto, A. G. A model of how the basal ganglia generate and use neural signals that predict reinforcement. in Models of information processing in the basal ganglia 249–270 (The MIT Press, 1995).

28. Rescorla, R. A. & Wagner, A. R. 3 A Theory of Pavlovian Conditioning: Variations in the Effectiveness of Reinforcement and Nonreinforcement. in (1972).

29. Schultz, W., Dayan, P. & Montague, P. R. A Neural Substrate of Prediction and Reward. Science 275, 1593–1599 (1997).

30. Kuo, S.-H. Ataxia. Contin. Minneap. Minn 25, 1036–1054 (2019).

31. Duncan, K., Semmler, A. & Shohamy, D. Modulating the Use of Multiple Memory Systems in Value-based Decisions with Contextual Novelty. J. Cogn. Neurosci. 1–13 (2019) doi:10.1162/jocn_a_01447.

32. Nicholas, J., Daw, N. D. & Shohamy, D. Uncertainty alters the balance between incremental learning and episodic memory. eLife 11, e81679 (2022).

33. Collins, A. G. E. & Frank, M. J. How much of reinforcement learning is working memory, not reinforcement learning? A behavioral, computational, and neurogenetic analysis. Eur. J. Neurosci. 35, 1024–1035 (2012).

34. Yoo, A. H. & Collins, A. G. E. How Working Memory and Reinforcement Learning Are Intertwined: A Cognitive, Neural, and Computational Perspective. J. Cogn. Neurosci. 34, 551–568 (2022).

35. Hoche, F., Guell, X., Vangel, M. G., Sherman, J. C. & Schmahmann, J. D. The cerebellar cognitive affective/Schmahmann syndrome scale. Brain 141, 248–270 (2018).

36. Chirino-Pérez, A. et al. Mapping the Cerebellar Cognitive Affective Syndrome in Patients with Chronic Cerebellar Strokes. The Cerebellum 21, 208–218 (2022).

37. McDougle, S. D. et al. Continuous manipulation of mental representations is compromised in cerebellar degeneration. Brain J. Neurol. awac072 (2022) doi:10.1093/brain/awac072.

38. Buckner, R. L. The cerebellum and cognitive function: 25 years of insight from anatomy and neuroimaging. Neuron 80, 807–815 (2013).

39. Koziol, L. F. et al. Consensus Paper: The Cerebellum’s Role in Movement and Cognition. Cerebellum Lond. Engl. 13, 151–177 (2014).

40. Alexander, M. P., Gillingham, S., Schweizer, T. & Stuss, D. T. Cognitive impairments due to focal cerebellar injuries in adults. Cortex J. Devoted Study Nerv. Syst. Behav. 48, 980–990 (2012).

41. Amokrane, N., Lin, C.-Y. R., Desai, N. A. & Kuo, S.-H. The Impact of Compulsivity and Impulsivity in Cerebellar Ataxia: A Case Series. Tremor Hyperkinetic Mov. 10, 43.

42. Buckner, R. L., Krienen, F. M., Castellanos, A., Diaz, J. C. & Yeo, B. T. T. The organization of the human cerebellum estimated by intrinsic functional connectivity. J. Neurophysiol. 106, 2322–2345 (2011).

43. Middleton, F. A. & Strick, P. L. Cerebellar Projections to the Prefrontal Cortex of the Primate. J. Neurosci. 21, 700–712 (2001).

44. Amokrane, N. et al. Impulsivity in Cerebellar Ataxias: Testing the Cerebellar Reward Hypothesis in Humans. Mov. Disord. 35, 1491–1493 (2020).

45. Chen, T. X. et al. Impulsivity Trait Profiles in Patients With Cerebellar Ataxia and Parkinson Disease. Neurology 99, e176–e186 (2022).

46. Thoma, P., Bellebaum, C., Koch, B., Schwarz, M. & Daum, I. The Cerebellum Is Involved in Reward-based Reversal Learning. The Cerebellum 7, 433 (2008).

47. Rustemeier, M., Koch, B., Schwarz, M. & Bellebaum, C. Processing of Positive and Negative Feedback in Patients with Cerebellar Lesions. Cerebellum Lond. Engl. 15, 425– 438 (2016).

48. McDougle, S. D. et al. Credit assignment in movement-dependent reinforcement learning. Proc. Natl. Acad. Sci. 113, 6797–6802 (2016).

49. Caligiore, D. et al. Consensus Paper: Towards a Systems-Level View of Cerebellar Function: the Interplay Between Cerebellum, Basal Ganglia, and Cortex. The Cerebellum 16, 203–229 (2017).

50. Hariri, A. R. The Emerging Importance of the Cerebellum in Broad Risk for Psychopathology. Neuron 102, 17–20 (2019).

51. Bellebaum, C. & Daum, I. Cerebellar involvement in executive control. The Cerebellum 6, 184–192 (2007).

52. Beuriat, P.-A. et al. A New Insight on the Role of the Cerebellum for Executive Functions and Emotion Processing in Adults. Front. Neurol. 11, (2020).

53. Mannarelli, D. et al. The Cerebellum Modulates Attention Network Functioning: Evidence from a Cerebellar Transcranial Direct Current Stimulation and Attention Network Test Study. The Cerebellum 18, 457–468 (2019).

54. Litman, L., Robinson, J. & Abberbock, T. TurkPrime.com: A versatile crowdsourcing data acquisition platform for the behavioral sciences. Behav. Res. Methods 49, 433–442 (2017).

55. Hoffman, M. D. & Gelman, A. The No-U-Turn Sampler: Adaptively Setting Path Lengths in Hamiltonian Monte Carlo. 31.

56. Team, S. D. Stan Reference Manual.

57. Vehtari, A., Gelman, A. & Gabry, J. Practical Bayesian model evaluation using leave-one-out cross-validation and WAIC. Stat. Comput. 27, 1413–1432 (2017).

58. Kalman, R. E. A New Approach to Linear Filtering and Prediction Problems. J. Basic Eng. 82, 35–45 (1960).

59. Nassar, M. R., Wilson, R. C., Heasly, B. & Gold, J. I. An Approximately Bayesian Delta-Rule Model Explains the Dynamics of Belief Updating in a Changing Environment. J. Neurosci. 30, 12366–12378 (2010).

